# CFPU: a cell-free processing unit for high-throughput, automated in vitro circuit characterization

**DOI:** 10.1101/2020.12.04.411470

**Authors:** Zoe Swank, Sebastian J. Maerkl

**Affiliations:** Institute of Bioengineering, School of Engineering, École Polytechnique Fédérale de Lausanne

## Abstract

Forward engineering synthetic circuits is at the core of synthetic biology. Automated solutions will be required to facilitate circuit design and implementation. Circuit design is increasingly being automated with design software, but innovations in experimental automation are lagging behind. Microfluidic technologies made it possible to perform *in vitro* transcription-translation (tx-tl) reactions with increasing throughput and sophistication, enabling screening and characterization of individual circuit elements and complete circuit designs. Here we developed an automated microfluidic cell-free processing unit (CFPU) that extends high-throughput screening capabilities to a continuous reaction environment, which is essential for the implementation and analysis of more complex and dynamic circuits. The CFPU contains 280 chemostats that can be individually programmed with DNA circuits. Each chemostat is periodically supplied with tx-tl reagents, giving rise to sustained, long-term steady state conditions. Using microfluidic pulse width modulation (PWM) the device is able to generate tx-tl reagent compositions in real-time. The device has higher throughput, lower reagent consumption, and overall higher functionality than current chemostat devices. We applied this technology to map transcription factor based repression under equilibrium conditions and implemented dynamic gene circuits switchable by small molecules. We expect the CFPU to help bridge the gap between circuit design and experimental automation for *in vitro* development of synthetic gene circuits.

## Introduction

Engineering gene regulatory networks from the bottom-up is a cornerstone of synthetic biology, providing a means to decipher natural biological systems [1, 2, 3] and develop new applications [4, 5, 6]. Developing new circuit designs *in vivo* requires design-build-test cycle iterations, where every iteration requires time-consuming molecular cloning steps. *In vitro* tx-tl systems on the other hand have proved to be powerful tools for accelerating the design-build-test cycle [3, 7, 8, 9, 10]. Recent technological advancements increased complexity and throughput of *in vitro* tx-tl experiments, such as the use of acoustic liquid handling robots [11, 12], or various microfluidic platforms [13, 14, 15]. Although these methods facilitated detailed studies of molecular circuits, they remain limited because of their use of simple batch reactions.

Continuous tx-tl reactions have been realized on microfluidic devices to enable *in vitro* implementation of dynamic circuits [16, 17]. These methods vastly expanded the type and complexity of networks that can be run in an *in vitro* environment compared to what is possible using batch reactions. However, these first generation chemostat devices are limited in throughput, creating a need for improved platforms that can carry out continuous tx-tl reactions at higher capacities. We developed a microfluidic device that combines the high-throughput capacities of batch reaction devices [15] and the sophistication of microfluidic chemostats [16, 17] to perform continuous tx-tl reactions in high-throughput.

In addition to being able to perform high-throughput, continuous tx-tl reactions, we developed a method to formulate reagent compositions on the fly to automatically screen condition space, or to dynamically perturb the system for analysis. Fluidic automation and process integration are increasingly recognized as being important capabilities for both chemical [18, 19] and biological methods [20, 21]. To enable fluidic process automation and dynamic, on-the-fly process changes we integrated a computer controlled fluidic metering strategy using microfluidic pulse width modulation (PWM) [22].

The resulting microfluidic CFPU enables the automated exploration of parameter space in 280 parallel running continuous reactions. Spotting and immobilizing DNA templates to program each chemostat allows screening of a large DNA sequence or concentration space. Furthermore, with the integration of computer controlled microfluidic PWM for on-chip metering and mixing of reaction components the platform can perform fully-automated and complex fluid processing tasks [22, 23]. By using different chemostat geometries, based on the design of Karzbrun *et al.* [17], we can alter reagent diffusion times. We integrated programmable DNA arrays, controlled DNA surface immobilization, highly-parallel chemostats, and programmable microfluidic PWM into a single CFPU and demonstrate its functionality by mapping the concentration and sequence space of transcription factor based repression at steady state and implemented a genetic toggle switch network.

## Results

### Design and characterization of the CFPU

The two-layer PDMS microfluidic device is fabricated using multi-layer soft-lithography [24] and consists of a control and a flow layer, whereby fluids on the flow layer can be manipulated by applying pressure to the control lines (Fig. 1A). The flow layer contains 280 unit cells, or chemostats, that can each run a unique and continuous tx-tl reaction (Fig. 1B). Each chemostat is connected to an exchange channel through which fresh reagents are flowed. The length of the connecting channel ranges from 50 *μ*m to 450 *μ*m, and a fixed width of 25 *μ*m. The flow layer inlets were optimized for mixing reagents by PWM [22]. The inlet flow channels are all of equal length as are the lengths of the control lines that regulate the the top five inlets, ensuring the PWM module accurately meters reagents. We incorporated a serpentine channel after the PWM module to allow reagent plugs formed by PWM to mix before reaching the beginning of the exchange channels.

**Figure 1:**
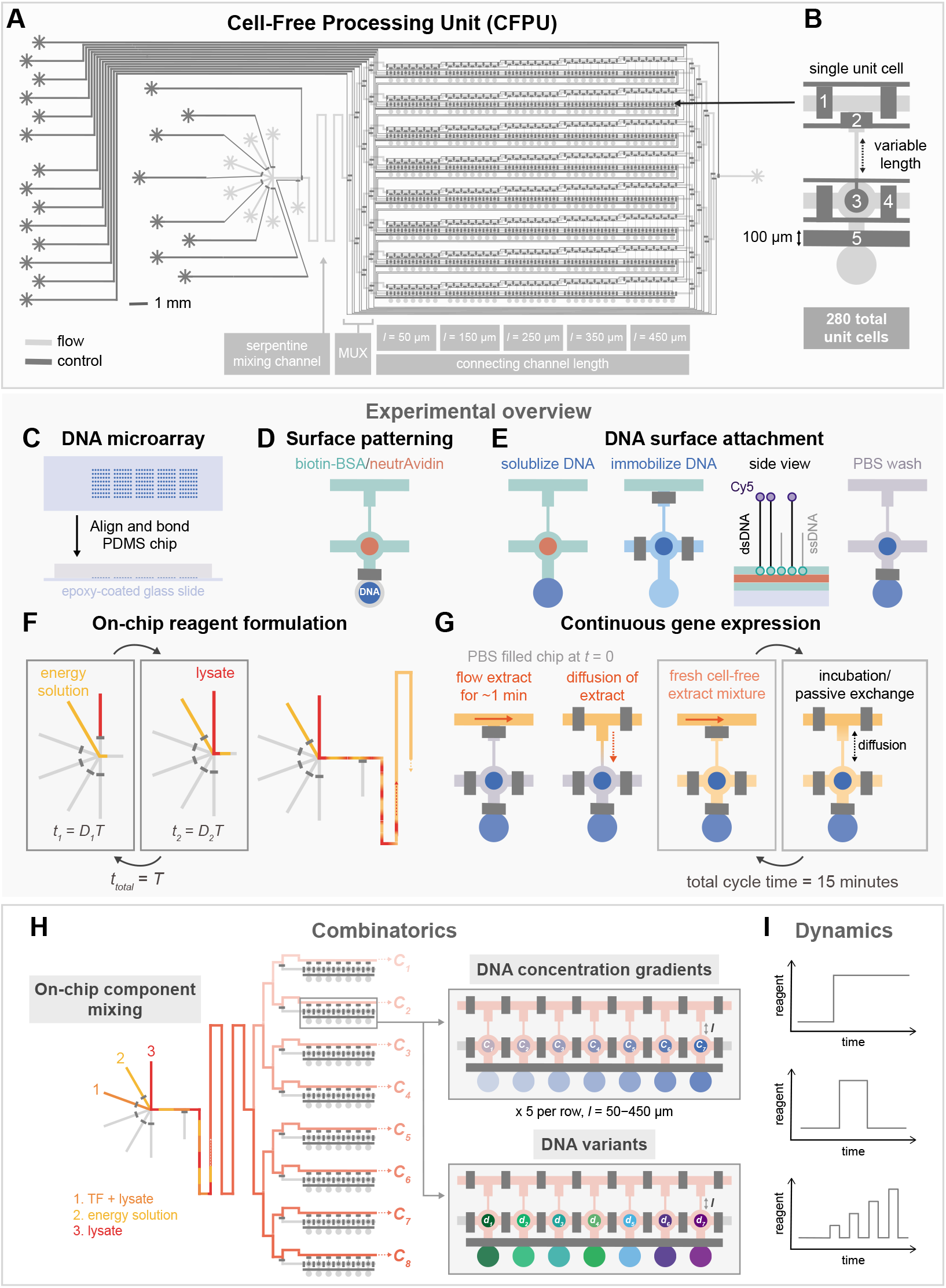
Overview of the CFPU architecture and functionality. **(A)** Illustration of the microfluidic CFPU design, highlighting the PWM optimized inlets, followed by the serpentine mixing channel and chemostat array with variable connecting channel lengths. **(B)** Design of a single unit cell. The pneumatic valves are described from top to bottom: 1) a pair of valves separates each unit cell in the exchange channel, 2) a valve prevents flow into the unit cell while the exchange channel is being replaced with fresh reagents, 3) the button valve enables surface patterning for DNA attachment, 4) sandwich valves separate the reaction chambers of each unit cell from one another, and 5) the neck valve isolates the DNA spot until surface patterning is complete. **C-G** A schematic summary of the experimental protocol: **(C)** DNA templates are spotted with a microarray robot onto an epoxy-coated glass slide, on top of which the PDMS chip is aligned. **(D)** The reaction chamber of the unit cell is patterned with biotin-BSA and neutrAvidin. **(E)** The DNA spot is solubilized, permitting the DNA to diffuse into the upper half of the unit cell and bind to the surface, after which all unbound DNA is washed away. **(F)** Lysate and energy solutions are then flowed and mixed on-chip with PWM. Each solution is flowed for a time *t* = *DT*, where *D* and *T* represent the duty cycle and cycle period, respectively. **(G)** Initially a cell-free extract solution is flowed onto the chip and diffuses into a unit cell that is filled with PBS. Every 15 minutes fresh cell-free extract is flowed through the exchange channel for ~1 minute followed by an incubation period that enables the exchange of reagents via diffusion. **(H)** Examples of combinatoric experiments that can be performed with this chip, including the generation of different concentrations of an input reagent using PWM. Each row of the device can be addressed with a specified reagent concentration, which is then combined with a range of DNA template concentrations or template variants, and different connecting channel lengths. **(I)** Reagent formulation is dynamic and a given reagent can be introduced into the device at user-defined times, allowing dynamic perturbations to be performed.

The device is prepared by first spotting fluorescently labeled and biotinylated linear DNA templates onto an epoxy-coated glass slide using a microarray robot (Fig. 1C). The PDMS chip is then aligned on top of the DNA microarray and bonded to the glass. After bonding, the upper half of each unit cell is patterned with biotin-BSA and neutrAvidin, resulting in a circular region of neutrAvidin coating (Fig. 1D), to which biotinlyated DNA can bind. Throughout this process the DNA spot remains isolated in the lower chamber, sealed off by the neck valve. Once surface patterning is completed, the DNA spot is solubilized, allowing the DNA to diffuse into the upper part of the unit cell and bind to the neutrAvidin coated area (Fig. 1E). Any unbound DNA is then washed from the device and tx-tl extract is introduced into the exchange channel by mixing the energy and lysate solutions directly on-chip with the PWM module (Fig. 1F). While the exchange channel is being replaced with fresh extract the bottom half of the unit cell is isolated with a valve that is then released to allow diffusion of fresh reagents into the unit cell. Another set of valves is actuated to isolate the unit cells from each other during incubation. The process of flowing fresh cell-free extract followed by an incubation step is repeated for up to 20 hours and achieves continuous, long-term gene expression (Fig. 1G). By periodically flowing fresh extract we are able to use far smaller reagent volumes (~3 μL/h), or approximately 20x less than previous methods (~60 μL/h) [17]. Additionally, on-chip mixing of lysate and energy components eliminates the need to cool the premixed reagents off-chip [25]. Using the multiplexing (MUX) valves (Fig. 1A), each row of the CFPU can be addressed with a specific tx-tl reagent composition (Fig. 1H). For instance, up to eight different concentrations of a transcription factor (TF) can be combined with the tx-tl components and tested in combination with different DNA template concentrations or DNA promoter variants. The added reagent, such as purified proteins or small molecules, are automatically formulated on-chip according to user defined time and concentration parameters, enabling dynamic gene circuit studies (Fig. 1I).

To confirm that the tx-tl extract solutions could be adequately mixed on-chip we tested various PWM cycle periods with duty cycles fixed at 50%. We used mCherry and GFP as tracers (~ 27 kDa) in the energy and lysate solutions, respectively, and monitored on-chip mixing at different locations downstream of the inlets. Directly after the inlets, the energy solution and lysate plugs can be clearly visualized for cycle periods ranging from 600 to 1400 ms (Fig. S1A). After passing through the serpentine channel, the two solutions have mixed for cycle periods less than 800 ms, and at the beginning of the exchange channel, mixing is complete for cycle periods up to 1200 ms long (Fig. S1B, C). As the CFPU architecture is separated into eight different rows, we also checked whether mixing is uniform across all rows and confirmed that for cycle periods of 600 and 1000 ms uniform mixing is achieved across all rows of the device (Fig. S1D, E).

Given that tx-tl solutions could be effectively mixed on-chip, we next tested continuous gene expression to characterize the effect of the different unit cell geometries (Fig. 2A). For a given DNA template concentration we observed a range of steady state expression levels depending on connecting channel length (Fig. 2B). This steady state can be disrupted by flowing PBS in place of the tx-tl extract mixture (Fig. S2). An increase in connecting channel length correlates with an increase in steady state GFP expression level as described previously [17]. Furthermore the time required to reach steady state levels increased for longer connecting channels. As described previously [15], binding of DNA templates on the surface of each unit cell and therefore their concentration can be precisely controlled by using a mixture of double stranded DNA templates and single stranded biotinlyated DNA oligos (Fig. 2C). When we varied the DNA template concentrations we observed a wide range of GFP expression levels as expected (Fig. 2D). Additionally, we achieved excellent chip to chip reproducibility (Fig. S3). Plotting GFP expression over time we saw that the higher the DNA template concentration or the longer the connecting channel the longer it took until a steady state was reached (Fig. S4). At lower DNA template concentrations, steady states can be reached after ~4-6 hours for all connecting channel lengths. For this reason, we performed all following experiments with a dsDNA to ssDNA ratio below 1:5.

**Figure 2:**
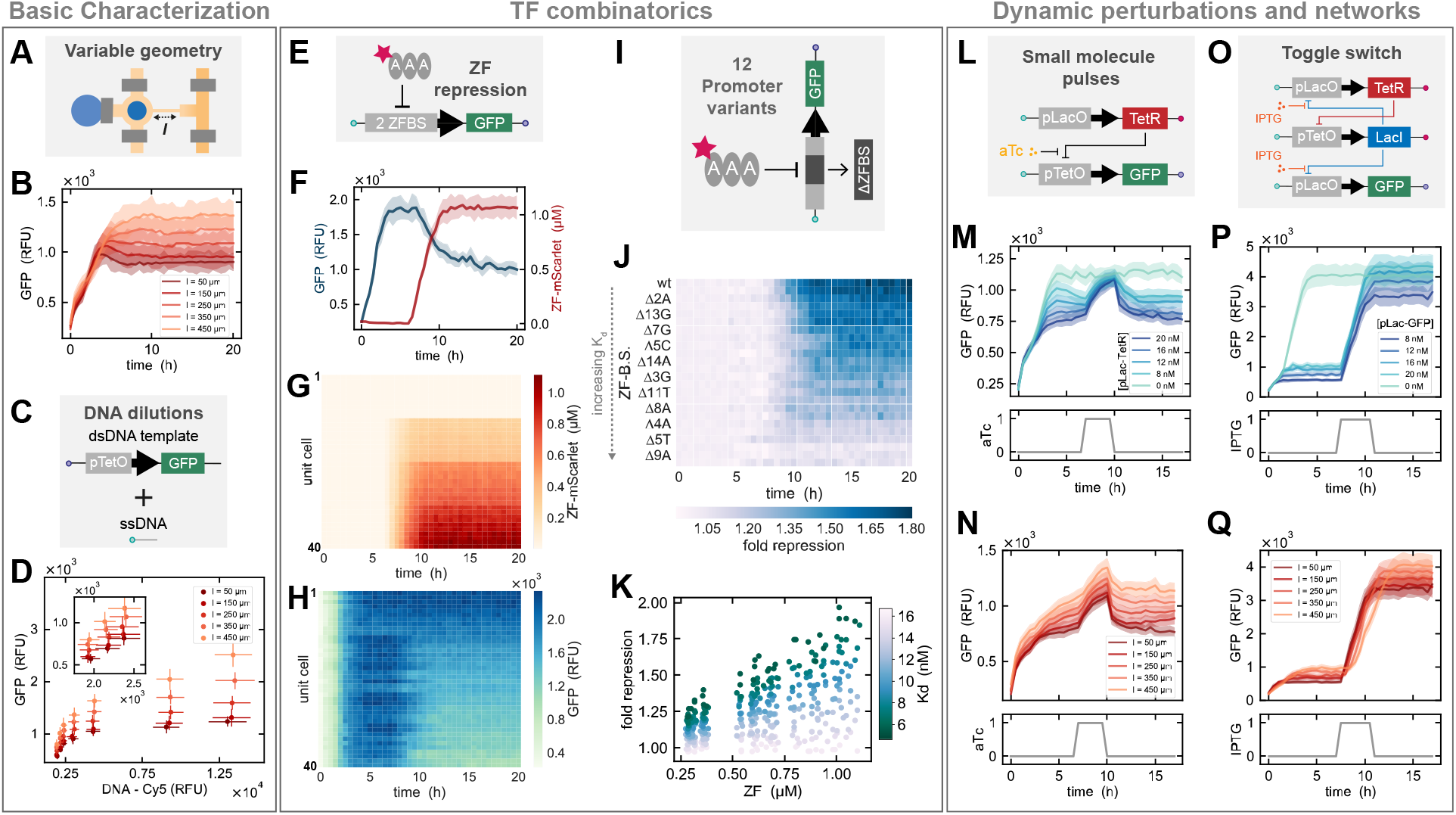
CFPU characterization and applications. **(A)** Protein expression using a single DNA template was carried out on-chip in order to characterize the effect of variable connecting channel lengths. **(B)** GFP expression versus time shown for a single DNA concentration and different connecting channel lengths. **(C)** DNA dilutions are made by mixing different ratios of dsDNA templates with short biotinylated ssDNA oligos. **(D)** The steady state GFP expression level versus the Cy5 fluorescence signal of dsDNA template attached to the surface for all connecting channel lengths and DNA dilutions. The inlaid plot shows the three lowest DNA concentrations. **(E)** PWM is used to mix three components on-chip to generate different concentrations of an mScarlet-tagged ZF which is then screened against a target template with two ZF binding sites in the promoter. **(F)** Example of time lapse traces for GFP expression and ZF-mScarlet signal. The lines represent the mean values ± SD (*n* = 2), where the SD is designated by the shaded regions. **(G)** ZF-mScarlet concentration over time for all unit cells containing a target template with two ZF binding sites in the promoter. **(H)** GFP expression versus time for the same unit cells shown in the preceding plot. **(I)** Multiple targets can be screened simultaneously on-chip, including promoter variants that contain mutated ZF binding sites. **(J)** Fold repression over time for targets containing modified ZF binding sites, shown in order of increasing *K_d_*, with the wild type binding site sequence at the top. The fold repression is calculated by dividing the signal obtained from no added ZF by the signal obtained from the highest quantity of added ZF (Duty cycle = 50 %). The time-lapse data shown in this plot corresponds to unit cells with a connecting channel length of *l* = 50 *μ*m. **(K)** All end-point fold repression values calculated for each duty cycle and connecting channel length, plotted according to the ZF binding site *K_d_*. Data shown in panels J and K was collected using two chips and represent the mean values ± SD (n = 2). **(L)** Illustration of the gene elements of the aTc controllable TetR repression assay. **(M)** GFP expression versus time for variable concentrations of pLac-TetR DNA template spotted inside unit cells with a connecting channel length of *l* = 50 *μ*m. **(N)** GFP expression profile over time shown for varying connecting channel lengths and a constant pLac-TetR DNA template concentration equal to 20 nM. **(O)** Sketch of the toggle switch gene circuit. **(P)** GFP expression versus time displayed for a range of pLac-GFP reporter concentrations spotted for unit cells with a connecting channel length of *l* = 50 *μ*m. The concentration of the TetR and LacI templates are kept constant, giving a range of repressor to reporter template ratios from 1:2 to 3:1. **(Q)** Time-lapse of GFP expression shown for different connecting channel lengths and a constant pLac-GFP reporter concentration equal to 8 nM. All data presented in M, N, P and Q represents mean values ± SD (*n* = 8).

### Mapping steady state repression by titrating repressor concentration

Forward engineering of gene regulatory networks often requires in depth characterization of the transcription factors used to regulate the network. For instance, repression using synthetic zinc finger (ZF) transcriptional regulators [26] can be tuned by varying both the concentration of ZF in the tx-tl reaction, as well as by tuning the ZF binding site [15]. As one proof-of-concept application, we screened the effect of different ZF-TF concentrations on repression at steady state.

We first tested whether it was possible to mix three different solutions with PWM: 1) a lysate solution containing a GFP tracer, 2) a plain lysate solution, and 3) an energy solution containing an mCherry tracer. As the ratio between the lysate and energy solution needs to be maintained at 1:1, the duty cycle of the energy solution is kept constant at 50%, while the remaining 50% of the duty cycle is divided between the two lysate solutions to achieve varying concentrations of GFP. For a cycle period of 1 s we observed a linear correlation between GFP concentration and duty cycle percentage (Fig. S5). At the same time the level of mCherry, which serves as a tracer in the energy solution, remained constant, indicating that the 1:1 ratio of energy and lysate solution was maintained.

After confirming that we could accurately mix three solutions on-chip with PWM, we designed an experiment to measure the steady state repression for *ZF_AAA_* at various concentrations in combination with a linear DNA template comprised of a GFP gene downstream of a λPR promoter, containing two ZF binding sites (Fig. 2E). We used a purified ZF-TF tagged with mScarlet, for on-chip visualization and quantification of protein concentration. PWM generated four ZF concentrations including a negative control and fed these into four clusters of rows using the MUX upstream of the exchange channels. Initially the tx-tl reaction was brought to steady state without ZF, and after 6.5 hours the different ZF concentrations were added to the respective sets of rows on the chip. As an example, we show the GFP signal and ZF-mScarlet concentration over time in two unit cells with the same connecting channel length and addressed with the same ZF input concentration (Fig. 2F). Looking at the ZF-mScarlet signal in each unit cell over time, we observe four distinct sections corresponding to the four ZF concentrations (Fig. 2G). However, within each quadrant we also notice a slight gradient, which is associated with the different connecting channel lengths. As the ZF diffuses into the unit cells we begin to see a decrease in GFP expression (Fig. 2H). We tested several promoter variants, containing one ZF binding site with different point mutations known to modulate repression to varying degrees (Fig. 2I). Overall four ZF concentrations were generated to test repression of 12 different DNA templates in five different unit cell geometries, for a total of 240 unique combinations, measured in replicates on two devices. The binding site mutations were chosen based on previously measured *K_d_*s [15], including two mutations that completely ablated repression (Δ9A, 5T) and nine mutations that covered the entire dynamic range between consensus and non-specific binding. If we consider a single unit cell dimension, we see that the fold repression for different target templates decreases over time as the *K_d_* associated with a given binding site increased (Fig. 2J). We observe a range of steady state fold repression values depending on the concentration of ZF within a given unit cell and a correlation between all fold repression values, ZF concentration, and the measured binding affinity for a given promoter (Fig. 2K). These results are in agreement with our previous batch analysis [15] and they provide a proof of concept that the microfluidic platform can be used to simultaneously screen target sequence space in conjunction with transcription factor concentrations in a steady state tx-tl reaction.

### Implementing dynamic gene circuits controlled by small molecules

After formulating different TFs concentrations on our chip at predesignated times, we went on to test whether it was possible to implement and control dynamic gene circuits. We set up a simple repression assay whereby TetR can repress a GFP encoding template that includes the Tet operator in its promoter region, unless a small molecule, anhydrotetracycline (aTc), is present and prevents TetR from binding to the operator (Fig. 2L). While holding the GFP DNA template concentration constant, we varied the concentration of TetR template and added a loading control template to maintain the same total DNA concentration. After allowing the reaction to reach steady state, we introduced aTc in a three hour pulse. As the concentration of TetR template was increased, we observed a corresponding decrease in the steady state expression levels of GFP (Fig. 2M). During the aTc pulse, repression was inhibited and GFP expression rose to the control level where no TetR template was present. Once aTc was no longer supplied to the unit cells, the GFP template was again repressed and the initial steady state re-established. Furthermore, we observed that the reaction time to the aTc pulse differs according to the connecting channel length (Fig. 2N).

As a final example we combined the ability to introduce small molecule inputs with dynamic gene circuits in order to implement and control a genetic toggle switch [27]. The toggle switch is composed of two linear DNA templates that encode two repressor proteins, TetR and LacI, where each protein encoding gene is downstream of mutually repressible promoters (Fig. 2O). A transient pulse of isopropyl-*β*-D-thiogalactopyranoside (IPTG) or aTc causes the circuit to switch from one state to another. Using different concentrations of a Lac-repressible GFP reporter we allowed the reaction to reach steady state, observing increasing levels of leak for higher concentrations of reporter template spotted. IPTG was introduced into the chip after 7 hours, causing TetR and GFP to be de-repressed (Fig. 2P). The toggle switch network remained stable in this state even after IPTG was not present after completion of the pulse. Similar to the varying response times we saw for aTc pulses before, the transition from one state to another occurs more slowly for longer connecting channel lengths, although we observed switching from a low to high state in all cases (Fig. 2Q). Considering that many genetic circuits incorporate small molecule control elements, we believe that our chip could serve as a useful platform for testing new network designs for which it is necessary to explore the parameter space of DNA template concentration together with small molecule regulators.

## Discussion

As synthetic biologists construct increasingly complex biomolecular circuits, the size of the parameter space governing these circuits becomes larger and more complex as well, requiring high-throughput and fast methods that can speed up the design-build-test cycle. Utilizing cell-free systems enables circuits to be optimized more rapidly than with traditional *in vivo* circuit construction and analysis since time-consuming molecular cloning steps can be bypassed. However, not all genetic networks can be tested in batch conditions and consequently need to be carried out in a system that mimics the dilutions which occur during cellular divisions. We therefore developed an automated CFPU that performs continuous *in vitro* tx-tl reactions in high-throughput, facilitating the characterization of biological parts and dynamic circuits. Each of the 280 on-chip chemostats can be programmed with a unique set of DNA templates. By precisely controlling the amount of DNA template attached to the surface of each unit cell, we can effectively scan a range of DNA concentrations that correlate with variable steady state protein expression levels. Different levels of protein expression can also be achieved by modifying the geometry of a given unit cell. On-chip reagent formulation by microfluidic PWM makes it possible to perform complex and fully-automated screens and generate complex reaction conditions in real-time. Combining the capacity to create custom reagent compositions with the potential to survey the DNA sequence and concentration space, the CFPU automates *in vitro* synthetic biology experiments.

## Materials and Methods

### Microfluidic chip fabrication

The designs for the flow and control layer of the device were drawn with AutoCAD software, we then used standard photolithography to fabricate the molds for each layer. SU-8 negative photoresist was used to create the control channel features (GM 1070, Gersteltec Sarl) with a height of 30 *μ*m, while AZ 9260 positive photoresist (Microchemicals GmbH) was used to generate flow channel features with a height of 15 *μ*m. Afterwards each of the wafers was treated with TMCS (trimethylchlorosilane) and coated with PDMS (polydimethylsiloxane, Sylgard 184, Dow Corning). For the control layer ~50 g of PDMS with an elastomer to crosslinker ratio of 5:1 was prepared, whereas for the flow layer a 20:1 ratio of elastomer to crosslinker was spin coated at 1800 rpm to yield a height of ~50 *μ*m. Both PDMS coated wafers were then partially cured for 20 minutes at 80 °C, after which devices from the control layer were cut out and the inlets for each control line were punched. Each control layer is then aligned onto the flow layer by hand using a Nikon stereo microscope. The aligned devices were then placed at 80 °C for 90 minutes, allowing the two layers to bond so that the entire device can be cut and removed from the flow wafer.

### Cell-free extract preparation

*E. coli* cell-free extract was prepared according to a published method [28] and a 4x energy solution was prepared based on the protocol described by Sun *et al.* [29]. As Kwon *et al.* screened a number of different parameters, we will briefly describe the protocol we have used. Cells were cultured with 2xYTP medium. After an initial overnight 5 mL culture, 1 mL of the overnight culture was added to a 500 mL Erlenmeyer flask with 200 mL of fresh medium. Four 200 mL cultures were incubated at 37 °C until an OD of ~2 at 600 nm was reached. Each culture was then separated into falcon tubes and centrifuged at 4 °C for 20 minutes at 4000 rpm. Cells were resuspended with a wash buffer (10 mM Tris, 14 mM magnesium glutamate, 60 mM potassium glutamate and 2 mM DTT) and centrifuged again at 4 °C for 10 minutes at 4000 rpm. Following three wash steps, all excess liquid is removed and the weight of each pellet is measured. The pellets are then flash frozen in liquid nitrogen and stored at −80 °C. The following day the cell pellets are thawed on ice and resuspended with 1 mL wash buffer per gram of cell pellet. Sonication was used to lyse the cells by applying a total energy input of ~400 J for a total cell suspension volume of ~1-1.5 mL. Cells were sonicated in an ice water bath with an amplitude of 50% with 10 s pulses interrupted by 10 s wait periods (Vibra-cell 75186 sonicator). Next the lysate solutions are centrifuged at 4 °C for 10 minutes at 12000 x g. The supernatent is removed and placed at 37 °C for 60 minutes to perform the run-off reaction. Lastly, the solution is centrifuged again at 4 °C for 10 minutes at 12000 x g and the supernatent is then aliquoted into 50 μL volumes, flash frozen and stored at −80 °C.

We prepared two different cell-free extracts originating from BL21 (DE3) and MC4100 cell strains. Lysates made from BL21 (DE3) cells were used for DNA dilution and ZF repression experiments, whereas MC4100 cell lysates were used for the small molecule inducible circuits as that strain is Lac- [30].

### Preparation of linear DNA templates

Linear DNA templates were generated by assembly PCR in order to link a downstream gene with a given promoter variant. The methods for generating the pλPR-GFP reporter templates used in the ZF repression assays have been previously described [15]. Gene specific PCR fragments were obtained for TetR and LacI from the repressilator plasmid [30] and combined thereafter with promoter PCR fragments that contained either the Lac or Tet operons, respectively. Using global primers we tagged each linear template at the 3’ end with biotin and at the 5’ end with either Cy3 or Cy5.

### Setting up high-throughput steady state cell-free tx-tl experiments

The process of DNA spotting, chip alignment and bonding, followed by chip priming, surface patterning and DNA immobilization has been described before [15]. Once the DNA templates had been immobilized at the surface of the unit cells, the button valve was actuated and the chip was washed with PBS for 15 minutes to ensure that any unbound DNA was removed and the DNA spot was visualized with fluorescence microscopy (Nikon Ti-SH-U). Before the experiment was started the button valve was released and the sandwich valves were closed.

A lysate solution was prepared by combining lysate with wash buffer in a 1:1 ratio and the energy solution was made up of 50% 4x energy solution, 2.5 *μ*M chi decoy DNA [31] or 3.5 *μ*M purified gamS [32] and water. The total volume of each solution should be between 30-40 μL in order to run the reaction for 15-20 hours. Each solution was drawn into a separate piece of FEP tubing (OD 1/16”, ID 1/32”, Upchurch) and connected to the microfluidic device inlet with a small piece of PEEK tubing (OD 1/32”, ID 0.18 mm, Vici). The pressure applied to the lysate and energy solution tubing was ~20 kPa during the experiment. A custom LabView program was written to control the pneumatic valves, the camera, and the microscope. Every 15 minutes a fresh mixture of cell-free extract was flowed through the exchange channel using a cycle period of 600 ms and a duty cycle of 50% for 100 total cycles (or ~60 s) to ensure that the entire channel volume has been replaced (Fig. S1F, G). To mix three components for the ZF repression experiment a cycle period of 1 s was used for a total of 60 cycles to result in the same overall flow time. Before the cell-free mixture was flowed, valve (1) was opened and valve (2) was closed, followed by a 10 s wait period to assure that valve (1) opened completely in all rows (Fig. 1A). Once the exchange channel was replaced with fresh reagents, valve (1) was closed and valve (2) was released permitting the diffusion of reaction components into and out of the unit cells. This process of exchange followed by incubation was carried out continuously for up to 20 hours while the chip was maintained in a temperature controlled box set to 33 °C. GFP expression was monitored in all unit cells over time by acquiring a fluorescence image every 30 minutes.

Switching from one cell-free mixture to another could be programmed in our LabView script. For instance, if a pulse of aTc or IPTG should be delivered to the unit cells at a given time then three solutions were prepared: 1) a lysate solution, 2) an energy solution and 3) an energy solution plus either 5 *μ*M of aTc or 250 *μ*M of IPTG. During the small molecule pulse the cell-free mixture will be generated from solutions (1) and (3), whereas at all other times it will be produced from solutions (1) and (2). For the ZF experiment three solutions were also used: 1) a lysate solution, 2) a lysate solution plus 5 *μ*M ZF-mScarlet and 3) an energy solution. Initially solutions (1) and (3) are used to reach steady state GFP expression levels, then all three solutions are used to generate different concentrations of ZF-mScarlet within the cell-free mixture. Furthermore the multiplexing valves are actuated accordingly to replace each exchange channel with a cell-free mixture containing a predefined ZF concentration. To ensure that the dead volume inside the serpentine mixing channel does not affect the next mixture generated, a PBS wash step is included every time the duty cycle percentage changes.

## Acknowledgments

We thank Ivan Istomin and Amir Shahein for providing us with the purified ZF-mScarlet protein. This work was supported by the European Research Council under the European Union’s Horizon 2020 research and innovation program Grant 723106 (SJM), and SNF Project grant 182019 (SJM).

## Author contributions

SZ, SJM developed the project. ZS performed experiments. ZS, SJM designed experiments, analyzed data, and wrote the manuscript.

## Supplementary Figures

**Figure S1:**
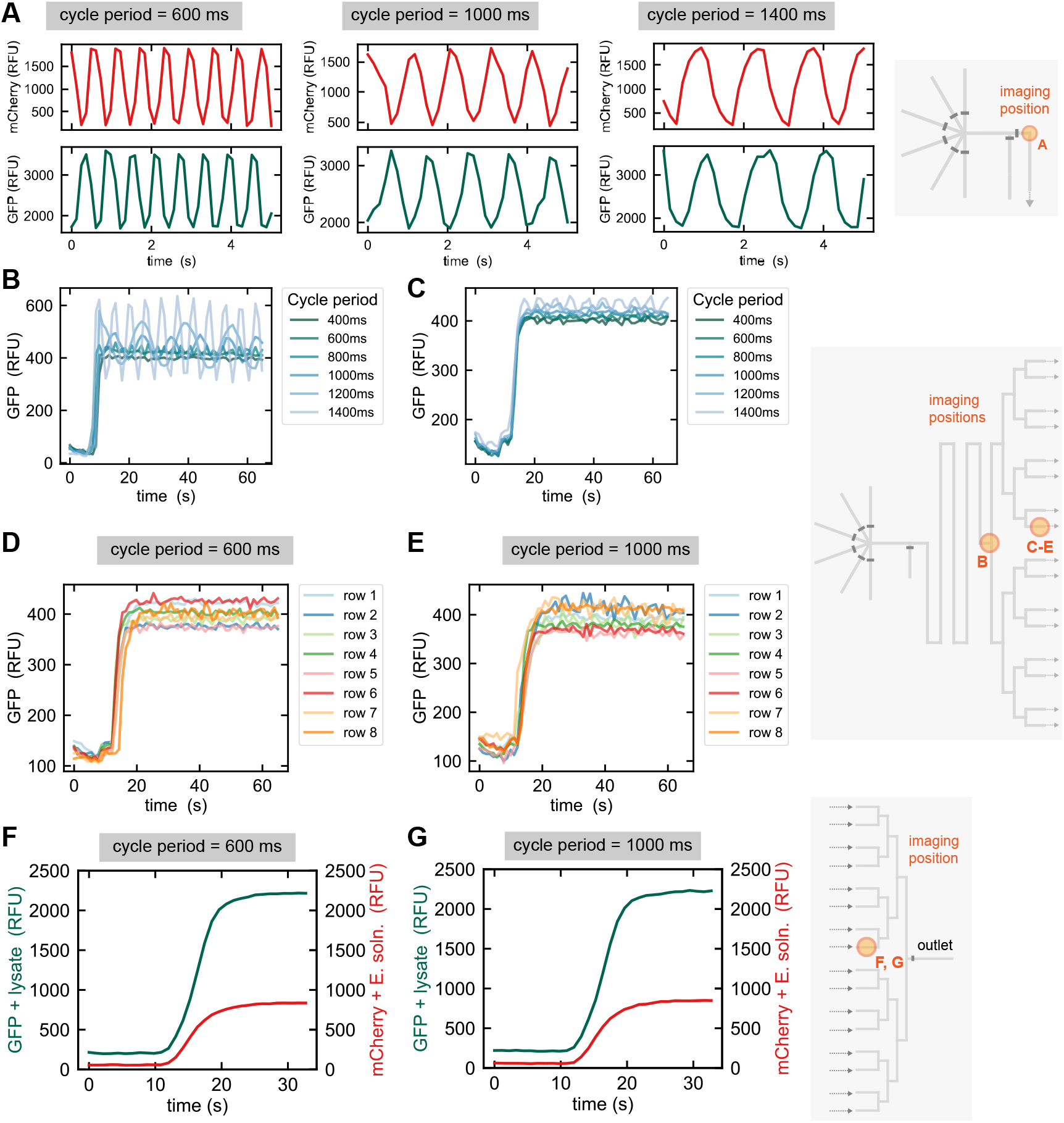
Characterization of PWM for on-chip mixing of lysate and energy solutions. **(A)** GFP (lysate) and mCherry (energy solution) signals versus time for cycle periods equal to 600, 1000 and 1400 ms. Images were acquired directly after the inlets, just before entering the serpentine channel. **(B, C)** GFP (lysate tracer) signal over time for a range of cycle periods as visualized at the end of the serpentine channel **(B)** and at the beginning of the exchange channel. **(D, E)** Time-lapse of the GFP (lysate tracer) signal acquired at the beginning of each exchange channel for cycle periods equal to 600 ms **(D)** and 1000 ms **(E)**. **(F, G)** GFP (lysate) and mCherry (energy solution) signals versus time for cycle periods equal to 600 ms **(F)** and 1000 ms **(G)** for images acquired at the end of the exchange channel. In all cases a duty cycle of 50% was maintained.

**Figure S2:**
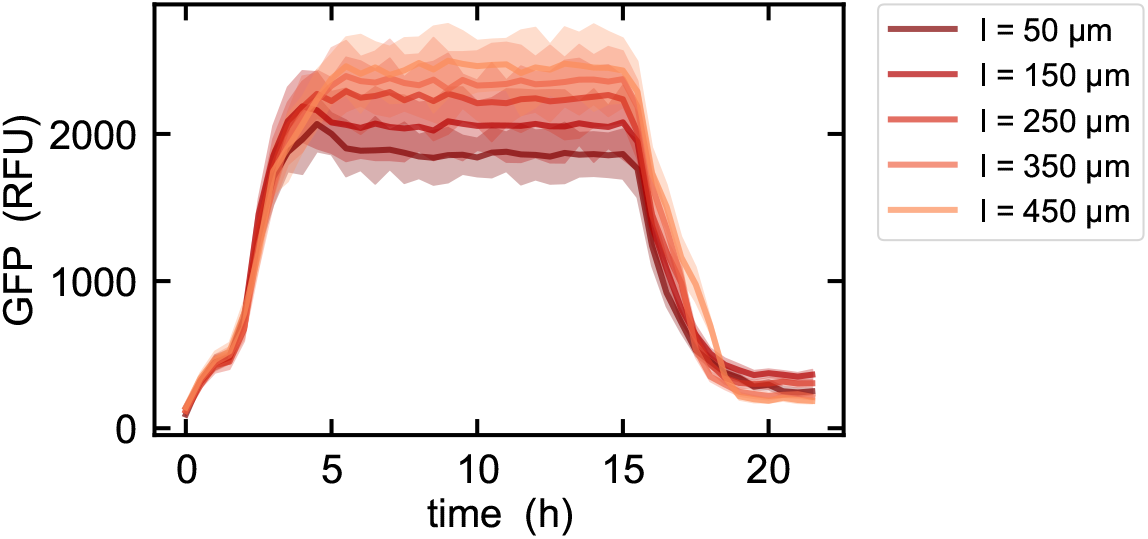
Disrupting steady state expression. Here we show the expression of GFP versus time for different unit cell connecting channel lengths. After 15 hours we stop flowing cell-free extract and instead flow PBS, showing that we can effectively terminate steady state GFP production in the absence of fresh reagents.

**Figure S3:**
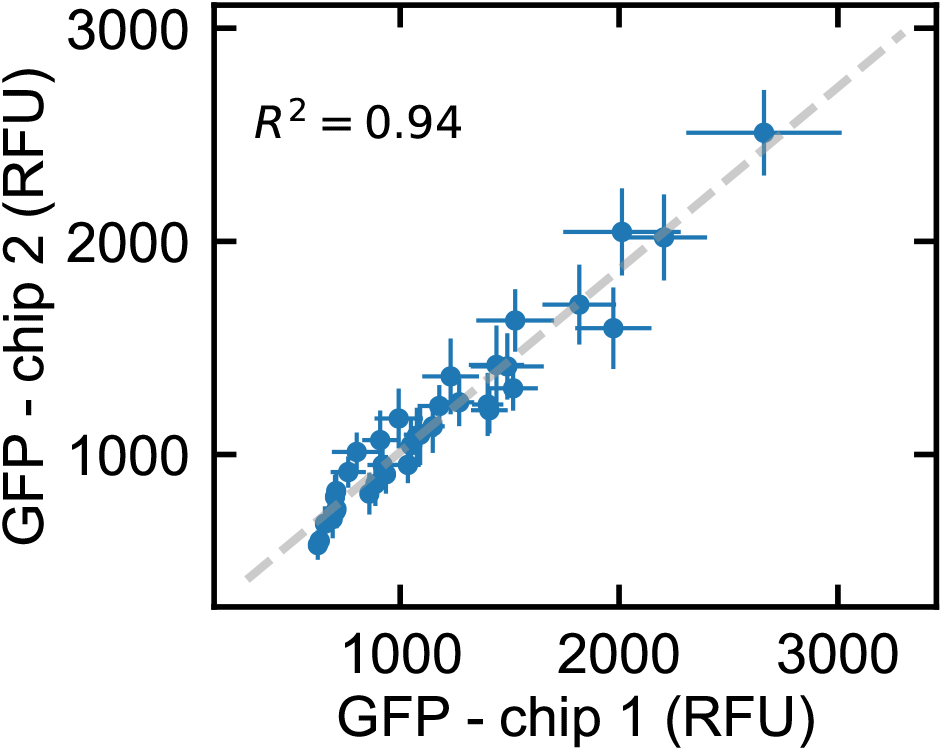
Chip-to-chip reproducibility. Steady state GFP expression values obtained for two separate chips for a range of DNA template dilutions. The same DNA microplate was used to spot both chips.

**Figure S4:**
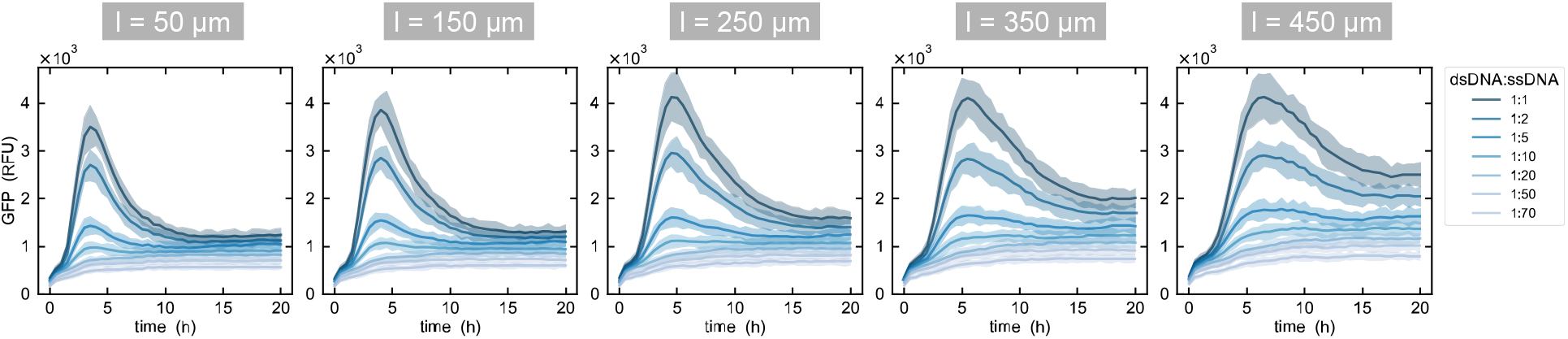
GFP expression for a range of DNA dilutions. GFP expression versus time for different ratios of dsDNA template to ssDNA oligo for each unit cell connecting channel length. All data presented represents mean values ± SD (n = 8)

**Figure S5:**
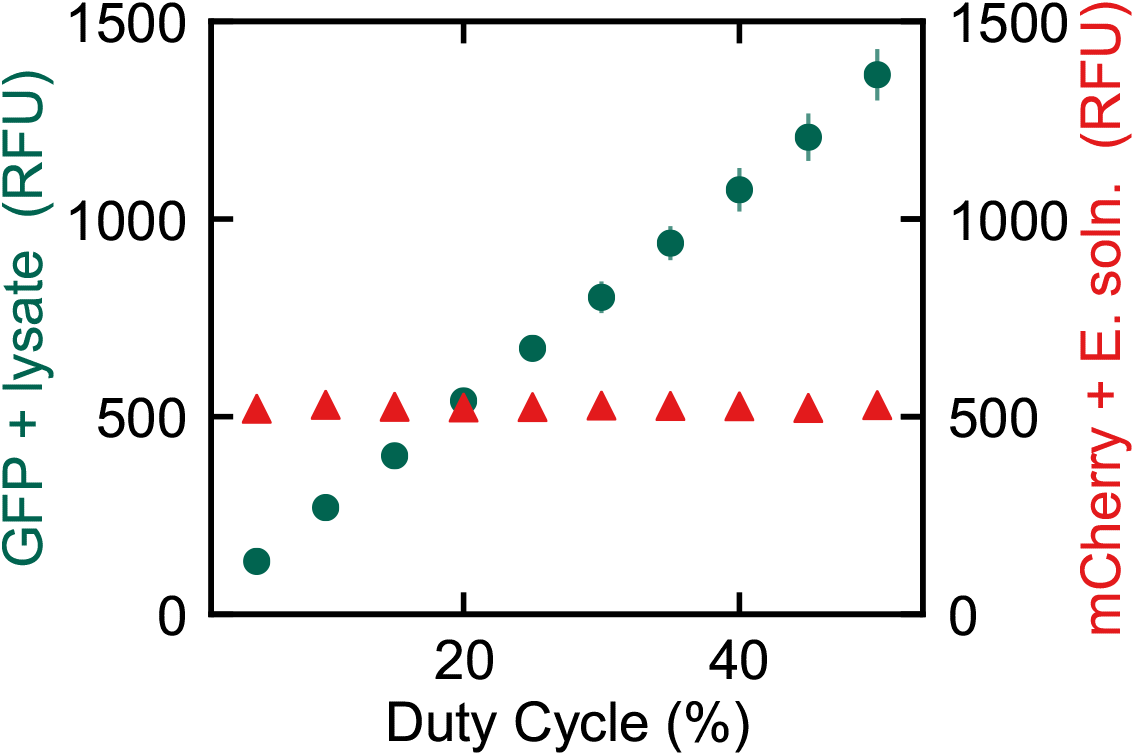
Mixing 3 components on-chip with PWM. Fluorescence intensity measured for GFP (lysate) and mCherry (energy solution) versus the duty cycle percentage corresponding to the lysate solution. The duty cycle for the energy solution was held constant at 50%.

## References

[1] Sprinzak, D. & Elowitz, M. B. Reconstruction of genetic circuits. Nature 438, 443–448 (2005).

[2] Bashor, C. J. & Collins, J. J. Understanding Biological Regulation Through Synthetic Biology. Annu. Rev. Biophys. 47, 399–423 (2018).

[3] Laohakunakorn, N. et al. Bottom-Up Construction of Complex Biomolecular Systems With Cell-Free Synthetic Biology. Front. Bioeng. Biotechnol. 8 (2020).

[4] Khalil, A. S. & Collins, J. J. Synthetic biology: applications come of age. Nature Reviews Genetics 11, 367–379 (2010).

[5] Cameron, D. E., Bashor, C. J. & Collins, J. J. A brief history of synthetic biology. Nat Rev Microbiol 12, 381–390 (2014).

[6] Silver, P. A., Way, J. C., Arnold, F. H. & Meyerowitz, J. T. Engineering explored. Nature 509, 166–167 (2014).

[7] Silverman, A. D., Karim, A. S. & Jewett, M. C. Cell-free gene expression: an expanded repertoire of applications. Nature Reviews Genetics 21, 151–170 (2020).

[8] Takahashi, M. K. et al. Rapidly Characterizing the Fast Dynamics of RNA Genetic Circuitry with Cell-Free Transcription–Translation (TX-TL) Systems. ACS Synth. Biol. 4, 503–515 (2015).

[9] Siegal-Gaskins, D., Tuza, Z. A., Kim, J., Noireaux, V. & Murray, R. M. Gene Circuit Performance Characterization and Resource Usage in a Cell-Free “Breadboard”. ACS Synth. Biol. 3, 416–425 (2014).

[10] Niederholtmeyer, H. et al. Rapid cell-free forward engineering of novel genetic ring oscillators. eLife 4, e09771 (2015).

[11] Moore, S. J. et al. Rapid acquisition and model-based analysis of cell-free transcription-translation reactions from nonmodel bacteria. PNAS 115, 4340–4349 (2018).

[12] Swaminathan, A., Hsiao, V. & Murray, R. M. Quantitative Modeling of Integrase Dynamics Using a Novel Python Toolbox for Parameter Inference in Synthetic Biology. bioRxiv (2017).

[13] Hori, Y., Kantak, C., Murray, R. M. & Abate, A. R. Cell-free extract based optimization of biomolecular circuits with droplet microfluidics. Lab Chip 17, 3037–3042 (2017).

[14] Fan, J. et al. Multi-dimensional studies of synthetic genetic promoters enabled by microfluidic impact printing. Lab Chip 17, 2198–2207 (2017).

[15] Swank, Z., Laohakunakorn, N. & Maerkl, S. J. Cell-free gene-regulatory network engineering with synthetic transcription factors. PNAS 116, 5892–5901 (2019).

[16] Niederholtmeyer, H., Stepanova, V. & Maerkl, S. J. Implementation of cell-free biological networks at steady state. PNAS 110, 15985–15990 (2013).

[17] Karzbrun, E., Tayar, A. M., Noireaux, V. & Bar-Ziv, R. H. Programmable on-chip DNA compartments as artificial cells. Science 345, 829–832 (2014).

[18] Adamo, A. et al. On-demand continuous-flow production of pharmaceuticals in a compact, reconfigurable system. Science 352, 61–67 (2016).

[19] Crowell, L. E. et al. On-demand manufacturing of clinical-quality biopharmaceuticals. Nature Biotechnology 36, 988–995 (2018).

[20] Boles, K. S. et al. Digital-to-biological converter for on-demand production of biologics. Nature Biotechnology 35, 672–675 (2017).

[21] Hartrampf, N. et al. Synthesis of proteins by automated flow chemistry. Science 368, 980–987 (2020).

[22] Woodruff, K. & Maerkl, S. J. Microfluidic Module for Real-Time Generation of Complex Multimolecule Temporal Concentration Profiles. Anal. Chem. 90, 696–701 (2018).

[23] Chang, J.-C., Swank, Z., Keiser, O., Maerkl, S. J. & Amstad, E. Microfluidic device for real-time formulation of reagents and their subsequent encapsulation into double emulsions. Scientific Reports 8, 8143 (2018).

[24] Thorsen, T., Maerkl, S. J. & Quake, S. R. Microfluidic Large-Scale Integration. Science 298, 580–584 (2002).

[25] Lavickova, B., Laohakunakorn, N. & Maerkl, S. J. A self-regenerating synthetic cell model. bioRxiv (2020).

[26] Blackburn, M. C., Petrova, E., Correia, B. E. & Maerkl, S. J. Integrating gene synthesis and microfluidic protein analysis for rapid protein engineering. Nucleic Acids Res 44, e68–e68 (2016).

[27] Gardner, T. S., Cantor, C. R. & Collins, J. J. Construction of a genetic toggle switch in Escherichia coli. Nature 403, 339–342 (2000).

[28] Kwon, Y.-C. & Jewett, M. C. High-throughput preparation methods of crude extract for robust cell-free protein synthesis. Scientific Reports 5, 8663 (2015).

[29] Sun, Z. Z. et al. Protocols for Implementing an Escherichia coli Based TX-TL Cell-Free Expression System for Synthetic Biology. JoVE (Journal of Visualized Experiments) e50762 (2013).

[30] Elowitz, M. B. & Leibler, S. A synthetic oscillatory network of transcriptional regulators. Nature 403, 335–338 (2000).

[31] Marshall, R., Maxwell, C. S., Collins, S. P., Beisel, C. L. & Noireaux, V. Short DNA containing *χ* sites enhances DNA stability and gene expression in E. coli cell-free transcription-translation systems. Biotechnol Bioeng 114, 2137–2141 (2017).

[32] Sun, Z. Z., Yeung, E., Hayes, C. A., Noireaux, V. & Murray, R. M. Linear DNA for Rapid Prototyping of Synthetic Biological Circuits in an Escherichia coli Based TX-TL Cell-Free System. ACS Synth. Biol. 3, 387–397 (2014).

